# Nociceptive stimuli activate the hypothalamus-habenula circuit to inhibit the mesolimbic reward system

**DOI:** 10.1101/2021.11.25.470073

**Authors:** Soo Min Lee, Yu Fan, Bonghyo Lee, Sang Chan Kim, Kyle B. Bills, Scott C. Steffensen, Hee Young Kim

## Abstract

Nociceptive signals interact with various regions of the brain, including those involved in physical sensation, reward, cognition, and emotion. Emerging evidence points to a role of nociception in the modulation of the mesolimbic reward system. The mechanism by which nociception affects dopamine (DA) signaling and reward is unclear. The lateral hypothalamus (LH) and the lateral habenula (LHb) receive somatosensory inputs and are structurally connected with the mesolimbic DA system. Here we show that the LH-LHb pathway is necessary for nociceptive modulation of this system. Our extracellular single-unit recordings and head-mounted microendoscopic calcium imaging revealed that nociceptive stimulation by tail-pinch excited LHb and LH neurons, which was inhibited by chemical lesion of the LH. Tail-pinch decreased extracellular DA release in the nucleus accumbens ventrolateral shell, which was blocked by disruption of the LH. Furthermore, tail-pinch attenuated cocaine-induced locomotor activity, 50-kHz ultrasonic vocalizations and reinstatement of cocaine-seeking behavior, which was inhibited by chemogenetic silencing of the LH-LHb pathway. Our findings suggest that nociceptive stimulation recruits the LH-LHb pathway to inhibit mesolimbic DA system and drug reinstatement.

## Introduction

Nociceptive stimuli include noxious pressure (e.g., tail-pinch), temperature (<10°C and >40°C), and chemicals (e.g., acids) [1]. The nociceptive signals are conveyed to the central nervous system from the periphery via spinal cord circuits and interact with many different brain areas, including those involved in physical sensation, reward, cognition, and emotion [2]. The mesolimbic dopamine (DA) system, sometimes referred to as the brain reward center, is a central nervous system circuit in which DA neurons in the ventral tegmental area (VTA) are connected to brain regions such as the nucleus accumbens (NAc), the prefrontal cortex, and the amygdala [3]. This system is critically involved in motivation, reward, and addiction [3]. Emerging evidence points to a role of nociception in the modulation of the mesolimbic system. Peripheral nerve injury, for instance, reduces morphine-induced conditioned place preference in mice and this effect is associated with DAergic activity in the NAc and the VTA [4]. In addition, nociceptive stimuli such as electric foot shocks and chemical injection during the self-administration training period strongly reduce drug-taking behavior such as fentanyl, cocaine, and methamphetamine in rodents [5–7]. Given the convergence of nociception with mesolimbic DA system, it is likely that nociception modifies the mesolimbic DAergic activity and influences drug reinstatement. However, to date, which neural circuit contributes to the delivery of nociceptive information to the mesolimbic DA system has not been fully known.

Nociceptive stimuli may require a series of neural circuits to arrive at the mesolimbic DA system [8]. The lateral habenula (LHb), an epithalamic structure, has been reported to be involved in processing information of peripheral sensation and nociceptive events and modulating motivational and cognitive processes [9]. Once activated, the LHb transmits glutamatergic projection to the rostromedial tegmental nucleus (RMTg) that projects GABAergic inputs to the VTA DA neurons, which eventually reduces DA release in the NAc [10, 11]. Studies have addressed the role of the LHb-RMTg-VTA connections in motivated behaviors and drug addiction [11–13]. The lateral hypothalamus (LH) has also been reported to process nociceptive signals [14, 15]. The LH and LHb are directly connected with each other by glutamatergic inputs arising from the LH [10]. Lazaridis et al. reported that the LH-LHb pathway encodes negative valence showing that optogenetic activation of the LHb-projecting LH glutamate neurons induces mice to switch from reward to non-reward in the probabilistic 2-choice switching task [16]. Given these findings, we hypothesized that the LH-LHb pathway conveys nociceptive signals to the mesolimbic DA system, thereby modulating cocaine-taking behavior and reinstatement of cocaine-seeking behavior.

To demonstrate this, we performed *in vivo* extracellular single-unit recording and calcium imaging to ascertain the effects of tail-pinch on neural activities of the LH and LHb and then chemically ablated the LH to examine whether the neural activity of the LHb is influenced by tail-pinch in the absence of the LH signals. We next recorded DA transients in the NAc ventrolateral shell during application of tail-pinch using *in vivo* fast-scan cyclic voltammetry (FSCV). We investigated effects of tail-pinch on behavioral changes induced by a single cocaine injection and cocaine-seeking/taking behaviors in the self-administration paradigm. Then, we investigated the role of the LH-LHb pathway in the effects of tail-pinch on cocaine-induced behavioral alterations by chemogenetic inhibition of the LH-LHb pathway.

## Results

### Excitation of the LHb and the LH by Tail-Pinch

To examine whether nociceptive stimulation excites the LH-LHb pathway, neural activities of the LHb and the LH following tail-pinch were investigated using *in vivo* extracellular single-unit recording or *in vivo* microendoscopic calcium imaging. Three different stimuli of brush, light pressure, and tail pinch were sequentially given to the tails for each 10 sec (Fig. 1A-C). While firing rates of the LHb neurons were not changed by brush and light pressure (8.5 g von Frey filament), application of tail-pinch evoked firing rates of the LHb neurons up to 182.25 ± 7.22% compared with the basal activity for 10 sec before tail-pinch (Fig. 1B and 1C; n=16 cells; one-way repeated ANOVA, F_(2,29)_=79.491).

**Figure 1.**
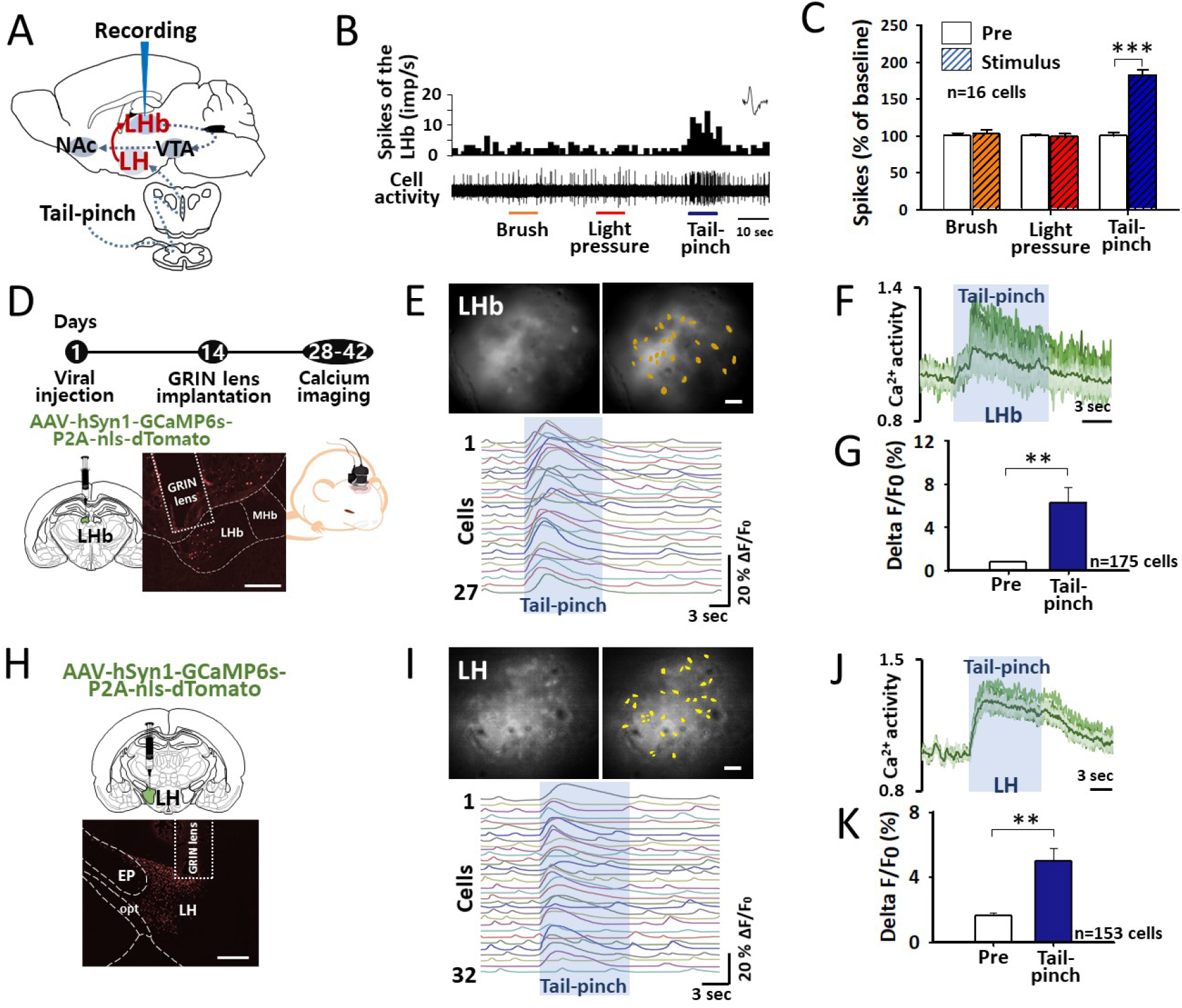
Effects of tail-pinch on the neural activities of the LHb and the LH. **(A)** A schematic diagram of *in vivo* extracellular single-unit recording in the LHb. **(B, C)** Neural activities of the LHb in responses to brush, light pressure, and tail-pinch for each 10 s. n=16 cells. ***p<0.001, Pre vs. Tail-pinch. **(D-G)** *in vivo* calcium imaging in the LHb following tail-pinch. GCaMP6s expression in the LHb and representative images of the GRIN lens track in the LHb (scale bar=4OO μm). An illustration of a rat with fluorescence microscope (D). A representative field of view in the LHb with ROIs (n=27 cells; scale bar=4O μm; top) and ΔF/F_0_ traces from the ROIs in the LHb (bottom; E). Normalized calcium transients of the ROIs in the LHb neurons (F). A bar graph of the averages of ΔF/F_0_ before and during tail-pinch for each 10 s in the LHb (n=6 rats; G; **p<0.005, Pre vs. Tail-pinch). **(H-K)** *in vivo* calcium imaging in the LH following tail-pinch. GCaMP6s expression in the LH (scale bar=4OO μm; H) A representative field of view in the LH with ROIs (n=32 cells; scale bar=4O μm; top) and ΔF/F_0_ traces from the ROIs in the LH (bottom; I). Normalized calcium transients of the ROIs in the LH neurons (J). A bar graph of the averages of ΔF/F_0_ before and during tail-pinch for each 10 s in the LH (n=5 rats; K; **p<0.009, Pre vs. Tail-pinch).

To confirm the excitatory effect of tail-pinch on the LHb neurons, *in vivo* calcium imaging was performed in the LHb (Fig. 1D-G). The rats with a calcium sensor GCaMP6s in the LHb were head-mounted with fluorescence microscope and then calcium transients following tail-pinch were recorded (Fig. 1D). When tail-pinch was applied for 10 sec, the calcium indicator GCaMP6s showed an initial rise followed by a sustained decrease in response to tail-pinch in the LHb neurons (Fig. 1E and 1F). The average of ΔF/F_0_ was 6.28 ± 1.43% during tail-pinch while the value before tail-pinch was 0.76 ± 0.09% (Fig. 1G; n=175 cells from 6 rats; paired *t-* test). We repeated this experiment in the LH neurons by using *in vivo* calcium imaging (Fig. 1H-K). The rats were given application of tail-pinch for 10 sec. Immediately after tail-pinch, fluorescence intensity of the calcium indicator GCaMP6s in the LH neurons markedly increased and declined steadily (Fig. 1I and 1J). The average of ΔF/F_0_ increased to 4.97±0.80% during tail-pinch from 1.61±0.21% (Fig. 1K; n=153 cells from 5 rats; paired *t*-test). These data indicate that nociceptive stimulation excites both the LHb and the LH neurons.

### The LH Mediation in Activation of the LHb by Tail-Pinch

To identify the mediation of the LH in transduction of nociceptive signals of tail-pinch to the LHb neurons, we ablated the LH by intracranial injection of neurotoxic ibotenic acid and measured a neural activity of the LHb following tail-pinch using *in vivo* extracellular single-unit recording or *in vivo* calcium imaging. Ibotenic acid was injected into the LH 7 days prior to the examinations (Fig. 2A and 2D). In *in vivo* extracellular single-unit recording, tail-pinch failed to induce excitation of the LHb neurons in the LH-lesioned rats (Fig. 2B and 2C; n=52 cells; one-way repeated ANOVA, F_(18,2)_=1.697). To perform *in vivo* calcium imaging, the LH-lesioned rats were injected with the calcium indicator GCaMP6s in the LHb and calcium-induced fluorescence changes following tail-pinch were monitored in the LHb (Fig. 2D). While tail-pinch induced a consistent increase in the fluorescence in intact rats (Fig. 1D), such increase was not observed in the LH-lesioned rats (Fig. 2E and 2F). In addition, the averages of ΔF/F_0_ before (Pre) and during tail-pinch (Tail-pinch) were 1.86 ± 0.46% and 2.37 ± 0.24%, respectively, but there was no significant difference between the two groups (n=121 cells from 4 rats; paired *t*-test). These data suggest that nociceptive signals are conveyed to the LHb via the LH.

**Figure 2.**
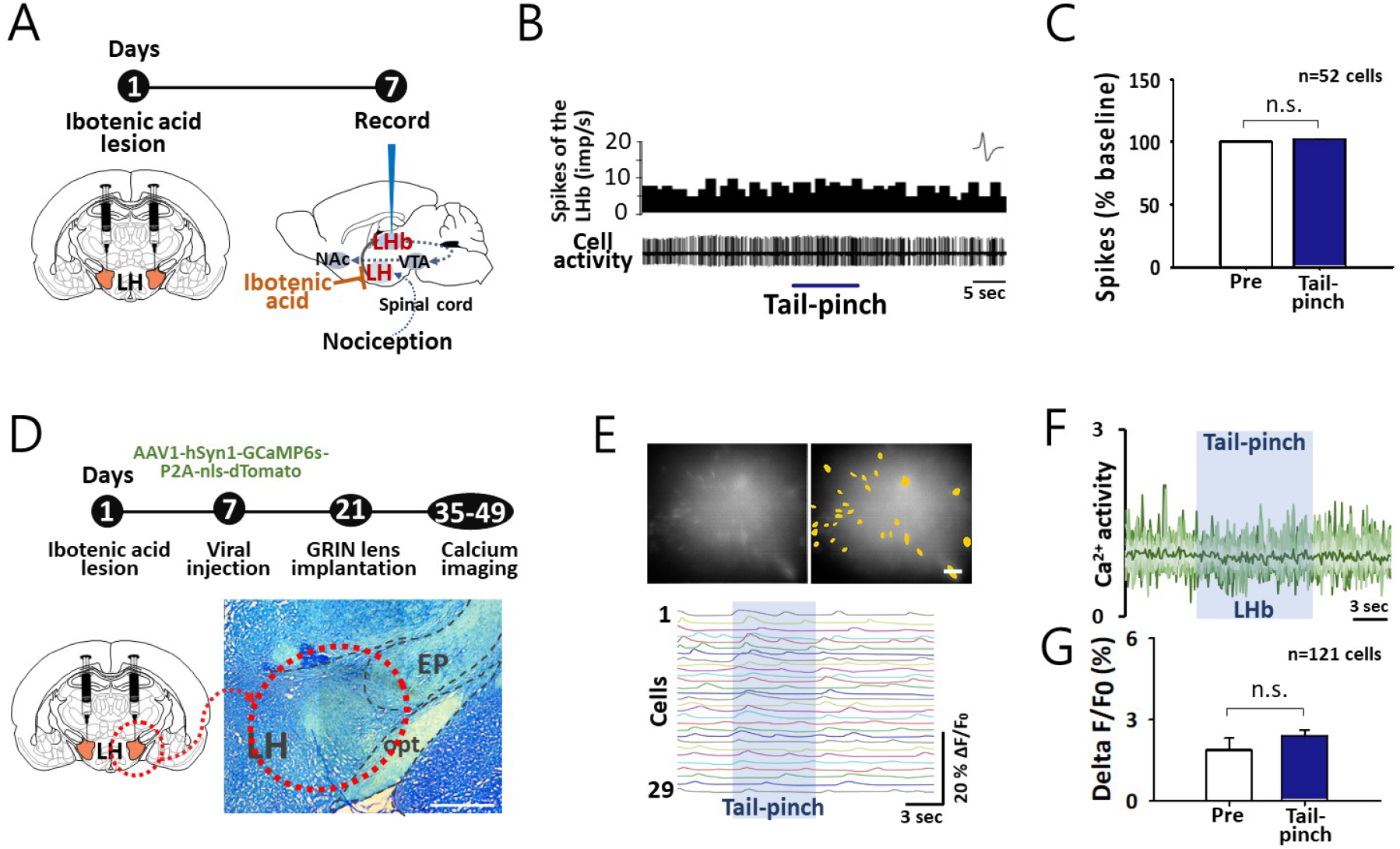
Effect of the LH lesion on the neuronal activity of the LHb following tail-pinch. **(A)** A timeline and diagrams of chemical lesion of the LH and *in vivo* extracellular single-unit recording in the LHb. **(B-C)** Neural activities of the LHb following tail-pinch in the rats with chemical lesion of the LH. n=52 cells. p=0.211. **(D)** A timeline of *in vivo* calcium imaging for neural activities of the LHb following tail-pinch in the rats with chemical lesion of the LH. A diagram and a representative image of chemical lesion of the LH (scale bar=400 μm). A diagram of GCaMP6s expression in the LHb. **(E-G)** A representative field of view in the LHb with ROIs (n=29 cells; scale bar=40 μm; top) and ΔF/F_0_ traces from the ROIs in the LHb (bottom; E). Normalized calcium transients of the ROIs in the LHb neurons (F). A bar graph of the averages of ΔF/F_0_ before and during tail-pinch for each 10 s in the LHb (n=4 rats; G). p=0.329.

### A Reduction of DA Release in the NAc Shell by Tail-Pinch and its Reversal by the LH Lesion

To determine whether nociceptive stimuli have an influence on mesolimbic DA release and if it was mediated via the LH, effects of tail-pinch on evoked DA release in the NAc ventrolateral shell were measured using *in vivo* FSCV in intact or the LH-lesioned rats. Ibotenic acid was injected into the LH a week prior to the experiments and then recordings were conducted in the ventrolateral part of the NAc shell (Fig. 3A and 3B). DA levels gradually decreased to 78.5 ± 4.85% 10 min after application of tail-pinch, but the decreased DA levels slowly recovered to the level of baseline when tail-pinch was removed (Fig. 3C; n=6 rats; one-way repeated ANOVA, F_(10,50)_=4.817). When tail-pinch was applied continuously for 30 min, DA levels steadily decreased to 63.9 ± 4.74% at the end of the stimulation compared to the values before tail-pinch (Fig. 3D and 3E; Tail-pinch group, n=7 rats) or Control (normal rats; n=6 rats). The sustained decrease of DA release by tail-pinch was not observed in the rats with chemical lesion of the LH (LH lesion+Tail-pinch group, n=8 rats; two-way repeated ANOVA, F_(28,140)_=3.943). These data suggest the mediation of the LH in nociceptive modulation of the mesolimbic DA system.

**Figure 3.**
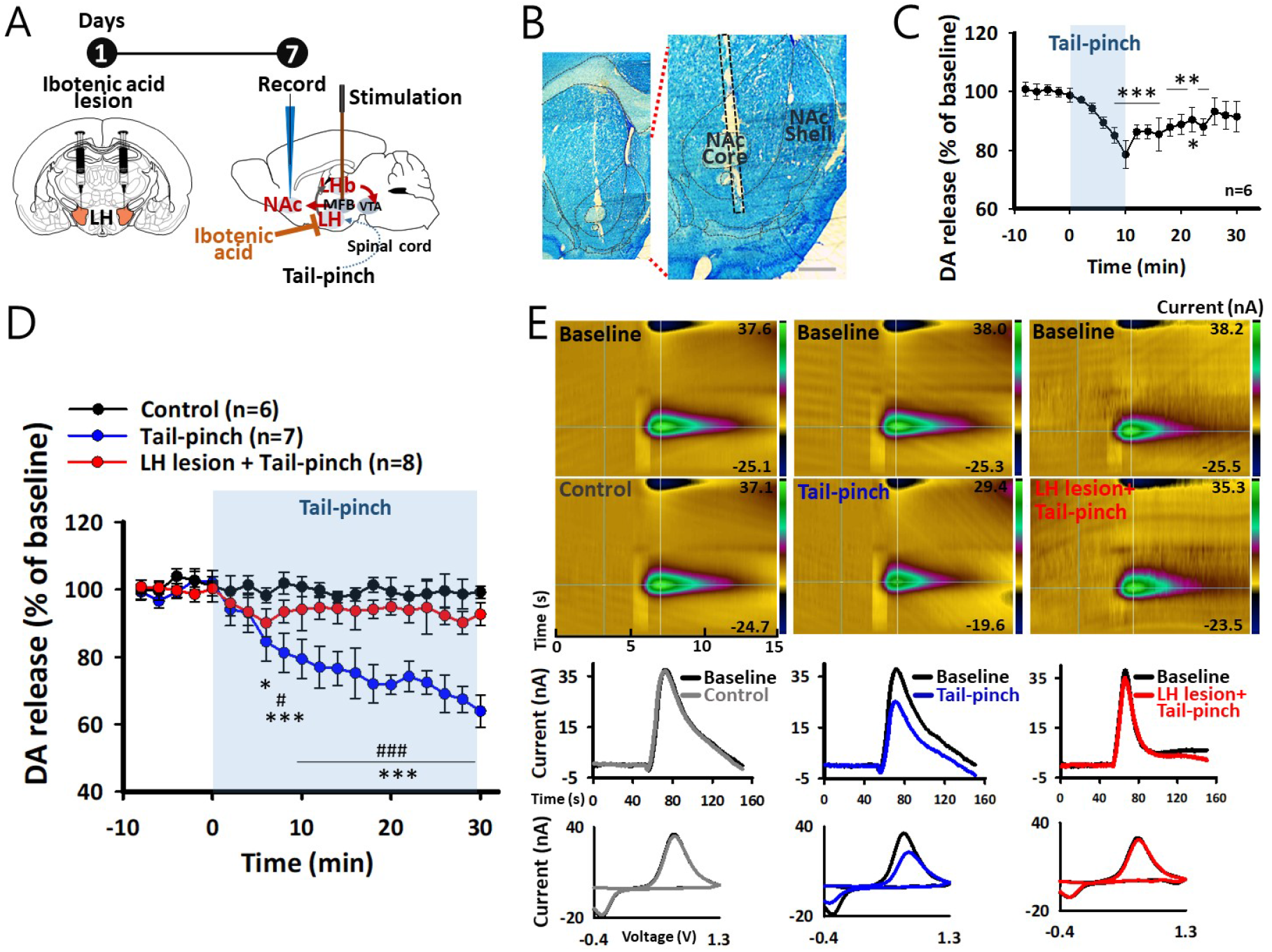
Effect of the LH lesion on the mesolimbic DA release following tail-pinch. **(A)** Schematics for *in vivo* FSCV in the rats with ibotenic acid lesion of the LH. Diagrams of ibotenic acid lesion of the LH and electrically evoked DA release in the NAc by stimulating the MFB. **(B)** Representative images of the recording site. A black dashed line indicates the track of a CFE. Bar=4OO μm. **(C)** Effect of tail-pinch for 10 min on the NAc DA release in naïve rats. n=6 rats. ***p<0.001, **p=0.002, 0.002, and 0.005, *p=0.03, before tail-pinch vs. after tail-pinch. **(D)** Comparison of the NAc DA release among control group (n=6 rats), Tail-pinch group (n=7 rats), and LH lesion+Tail-pinch group (n=8 rats). *p=0.012, ***p<0.001, Tail-pinch vs. Control; #p=0.016, ###p<0.001, Tail-pinch vs. LH lesion+Tail-pinch. **(E)** Representative pseudo-color plots with color bars indicating the current. Time-series plots indicate the current vs. time traces for DA release in each group. Each cyclic voltammogram corresponds to the above pseudo-color plots.

### Suppression of Cocaine-Induced Psychomotor Activities by Tail-Pinch and Mediation of the LH-LHb Pathway

On the basis of our electrophysiological and microendoscopic findings that tail-pinch activated the LH-LHb pathway and thus suppressed extracellular DA release in the NAc ventrolateral shell, we further explored the effects of tail-pinch on acute cocaine-enhanced locomotor activity and emission of 50-kHz USVs, known to be associated with mesolimbic DA levels [17]. Locomotor activity and 50-kHz USVs were recorded in custom-built chambers simultaneously (Fig. 4A). Cocaine administration rapidly increased locomotion with a peak at 10 min, followed by a steady decrease over the 60 min (Fig. 4B; Coc. group, n=6 rats). Tail-pinch significantly inhibited the cocaine-enhanced locomotion (Coc.+Tail-pinch group, n=6 rats), compared to the Coc. group, while tail-pinch itself did not affect locomotor activity in normal rats (Tail-pinch group, n=6 rats; two-way repeated ANOVA, F_(10,50)_=31.814). Furthermore, we analyzed the number of 50-kHz USVs, detected during the locomotor activity (Fig. 4C). Cocaine administration evoked a large number of 50-kHz USVs, compared to the basal value before cocaine injection, which was strongly reduced by application of tail-pinch (two-way ANOVA, F_(2,24)_=64.629). These data suggest a reversal of the cocaine-induced psychomotor activities by nociception.

**Figure 4.**
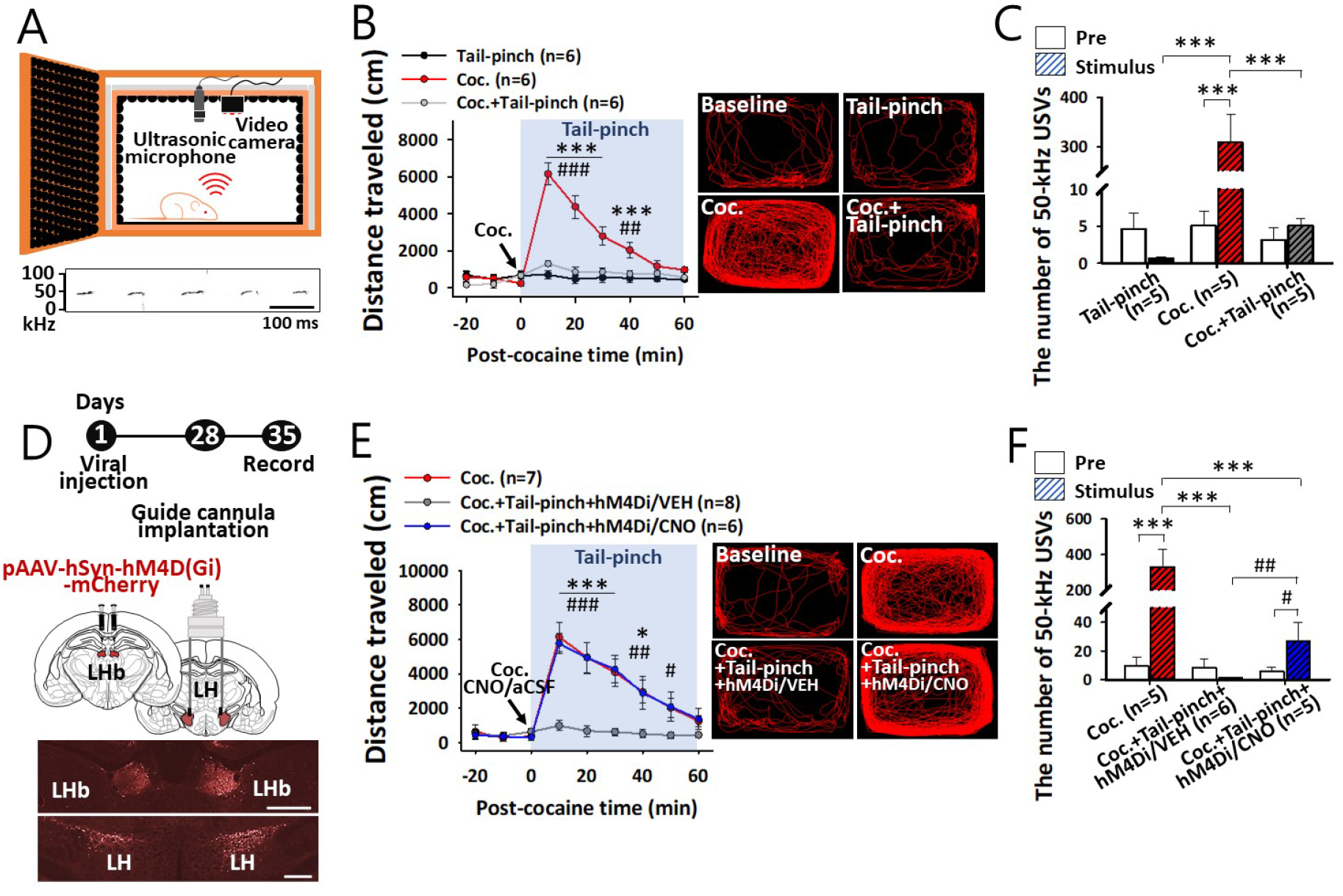
Effect of chemogenetic silencing of an LH-LHb pathway on tail-pinch inhibition of cocaine-induced locomotion and emission of 50-kHz USVs. **(A)** Illustration of a freely moving rat in a customized USV chamber (top) and a representative spectrogram of 50-kHz USVs (bottom). **(B, C)** Effect of tail-pinch on cocaine-enhanced locomotion and 50-kHz USVs in rats. Effect of tail-pinch on cocaine-induced locomotion in rats (left of B; ***p<0.001, Coc. vs. Tail-pinch; ##p=0.005, ###p<0.001, Coc. vs. Coc.+Tail-pinch; n=6/group). Representative locomotion tracks for 30 min after tail-pinch and/or cocaine injection (right of B). Average of 50-kHz USVs for 30 min before and after tail-pinch and/or cocaine injection (C; ***p<0.001, Coc. vs. Pre, Tail-pinch, and Coc.+Tail-pinch; Tail-pinch, n=5; Coc., n=5; Coc.+Tail-pinch, n=5). Tail-pinch, tail-pinch in naïve rats; Coc., cocaine injection only; Coc.+Tail-pinch, tail-pinch in cocaine-treated rats. **(D)** A timeline and diagrams of hM4Di injection in the LHb and guide cannula implantation in the LH. Representative images of hM4Di expression in the LHb and the LH. Bar=400 μm. **(E, F)** Effect of chemogenetic silencing of the LH-LHb pathway on tail-pinch inhibition of cocaine-induced locomotion and emission of 50-kHz USVs. Effect of chemogenetic silencing of the LH-LHb pathway on tail-pinch inhibition of cocaine-induced locomotion (left of E; *p=0.011, ***p<0.001, Coc.+Tail-pinch+hM4Di/VEH vs. Coc.; #p=0.039, ##p=0.005, ###p<0.001, Coc.+Tail-pinch+hM4Di/VEH vs. Coc.+Tail-pinch+hM4Di/CNO; Coc., n=7; Coc.+Tail-pinch+hM4Di/VEH, n=8; Coc.+Tail-pinch+hM4Di/CNO, n=6) and representative locomotion tracks for 30 min after tail-pinch, cocaine injection, and/or either CNO or aCSF infusion (right of E). A retrograde viral vector encoding an inhibitory DREADD (hM4Di) was injected into the LHb and CNO was intracranially infused into the LH. Average of 50-kHz USVs for 30 min before and after tail-pinch, cocaine injection, and/or either CNO or aCSF infusion (F; ***p<0.001, Coc. vs. Pre, Coc.+Tail-pinch+hM4Di/VEH, and Coc.+Tail-pinch+hM4Di/CNO; #p=0.011, ##p=0.008, Coc.+Tail-pinch+hM4Di/CNO vs. Pre and Coc.+Tail-pinch+hM4Di/VEH; Coc., n=5; Coc.+Tail-pinch+hM4Di/VEH, n=6; Coc.+Tail-pinch+hM4Di/CNO, n=5). Coc., cocaine injection only in hM4Di-expressed rats; Coc.+Tail-pinch+hM4Di/VEH, cocaine injection, tail-pinch and aCSF infusion into the LH in hM4Di-expressed rats; Coc.+Tail-pinch+hM4Di/CNO, cocaine injection, tail-pinch and CNO infusion into the LH in hM4Di-expressed rats.

To evaluate mediation of the LH-LHb pathway in the inhibitory effects of tail-pinch on the cocaine-induced psychomotor activities, a retrograde viral vector encoding an inhibitory DREADD (hM4Di) was injected into the LHb, CNO was intracranially infused into the LH (LH-LHb:hM4Di/CNO) in order to silence the LH-LHb pathway, and then effects of tail-pinch on cocaine-induced behaviors were explored (Fig. 4D). As shown in Fig. 4E, cocaine administration enhanced locomotor activity (Coc. group, n=7 rats), which was suppressed by tail-pinch in the rats with LH-LHb:hM4Di/VEH (Coc.+Tail-pinch+hM4Di/VEH group, n=8 rats). The inhibitory effects of tail-pinch on the cocaine-induced locomotion were almost completely blocked by intracranial CNO infusion in the rats with LH-LHb:hM4Di (Coc.+Tail-pinch+hM4Di/CNO group, n=6 rats; two-way repeated ANOVA, F_(10,50)_=16.004). These effects were further confirmed by measuring the number of 50-kHz USVs (Fig. 4F). The increased emission of 50-kHz USVs by cocaine injection was suppressed by tail-pinch stimulation in the rats with LH-LHb:hM4Di/VEH (Coc.+Tail-pinch+hM4Di/VEH group). In contrast, in the rats with LH-LHb:hM4Di, intracranial CNO infusion significantly alleviated the inhibitory effects of tail-pinch on cocaine-induced emission of 50-kHz USVs, compared to aCSF infusion (two-way ANOVA, F_(2,26)_=207.049). These results indicate that the inhibitory effects of tail-pinch on the cocaine-induced hyperlocomotion and positive affective states (50-kHz USVs) are mediated via the LH-LHb pathway.

### Attenuation of Cocaine-Taking/Seeking Behaviors by Tail-Pinch and Mediation of the LH-LHb Pathway

To further examine whether tail-pinch could suppress cocaine-taking behavior and reinstatement of cocaine-seeking behavior, we employed the cocaine self-administration paradigm (Fig. 5A and 5B). Mediation of the LH-LHb pathway was investigated by using chemogenetic inhibition (LH-LHb:hM4Di/CNO). Initially, effects of tail-pinch on a natural reward were investigated using food training (Fig. 5C; one-way ANOVA, F_(3,32)_=1.000). Application of tail-pinch throughout the test session (3 h; Tail-pinch session) did not affect consumption of food pellets. During the cocaine self-administration training (Fig. 5D), rats were trained to intravenous cocaine infusions (0.5 mg/kg/infusion; Coc. group, n=6 rats). Application of tail-pinch during the cocaine self-administration training inhibited the establishment of cocaine self-administration (Coc.+Tail-pinch group, n=5 rats; two-way repeated ANOVA, F_(9,36)_=1.355). As shown in Fig. 5E, after rats were trained for establishing the cocaine self-administration (Coc. group, n=5 rats; Coc.+Tail-pinch group, n=6 rats; Coc.+Tail-pinch+hM4Di/VEH group, n=5 rats; Coc.+Tail-pinch+hM4Di/CNO group, n=6 rats), we tested the effects of tail-pinch on cocaine intakes (Test 1; Fig. 5E-G; 5F, one-way ANOVA, F_(3,18)_=0.00374; 5G, one-way ANOVA, F_(3,18)_=1.275). Next, the rats were trained for extinguishing the cocaine self-administration and we investigated the effects of tail-pinch on reinstatement of cocaine-seeking behavior (Test 2; Fig. 5H-J). In the test 1, although tail-pinch tended to reduce the number of cocaine infusions, there were no statistically significant differences among the groups in the number of cocaine intakes as well as active/inactive lever responses (Fig. 5E-G). The rats were then subjected to extinction sessions which the cocaine solution was replaced with saline and the test 2 was conducted (Fig. 5H-J). While a single priming injection of cocaine (0.5 mg/kg/mL) produced robust active lever responses that indicate reinstatement of cocaine-seeking behavior (Coc. group; Fig. 5I), tail-pinch suppressed the cocaine-primed active lever responses (one-way ANOVA, F_(3,17)_=7.190). The inhibitory effect of tail-pinch on the reinstatement of cocaine-seeking behavior was suppressed by pretreatment of intracranial CNO infusion in the rats expressing hM4Di in the LH-LHb pathway. However, tail-pinch did not affect the number of cocaine intakes and inactive lever responses (Fig. 5H and 5J; 5J, one-way ANOVA, F_(2,14)_=1.130). These data indicate that nociceptive stimulation attenuated the cocaine-taking/seeking behaviors via the LH-LHb pathway.

**Figure 5.**
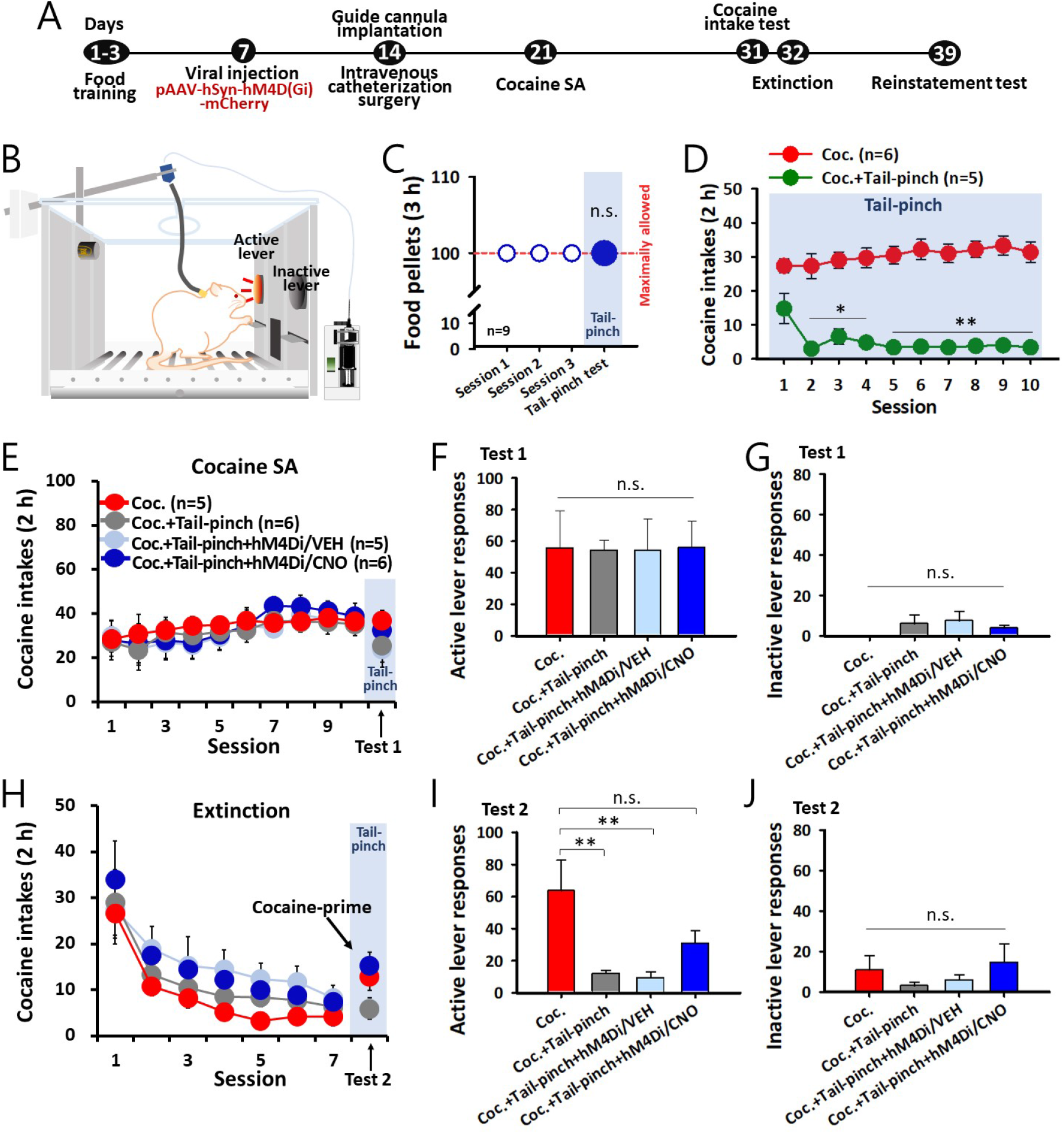
Effects of tail-pinch on cocaine-taking/seeking behaviors and its reversal by chemogenetic silencing of the LH-LHb pathway. **(A, B)** Experimental procedures for the cocaine self-administration. **(C)** Effect of tail-pinch on the consumption of food pellets during food training session. The total number of food pellets was limited to 100 in each experiment. n=9 rats. p=1.000. **(D)** Effect of tail-pinch on cocaine intakes during cocaine self-administration training. Coc. (cocaine self-administration training; n=6); Coc.+Tail-pinch (tail-pinch during cocaine self-administration training, n=5). *p=0.034, 0.024, and 0.012, **p=0.008, 0.005, 0.007, 0.006, 0.004, and 0.006, Coc. vs. Coc.+Tail-pinch. **(E-G)** Effect of tail-pinch on cocaine intakes after acquisition of cocaine self-administration over 10 sessions. Time courses of cocaine infusions (E). The numbers of active lever responding following tail-pinch (F; p=1.000). The number of inactive lever responding following tail-pinch (G; p=0.313). (**H-J**) Inhibition by tail-pinch of cocaine-primed reinstatement of cocaine-seeking behavior and its reversal by chemogenetic silencing of the LH-LHb pathway. Time courses of cocaine infusions (H). The numbers of active lever responding following tail-pinch (I; **p=0.006 and 0.004, Coc. vs. Coc.+Tail-pinch and Coc.+Tail-pinch+hM4Di/VEH). The number of inactive lever responding (J; p=0.351). Coc., cocaine priming injection only (n=5); Coc.+Tail-pinch, cocaine priming injection + tail-pinch; Coc.+Tail-pinch+hM4Di/VEH, cocaine priming injection, tail-pinch and aCSF infusion into the LH in hM4Di-expressed rats (n=5); Coc.+Tail-pinch+hM4Di/CNO, cocaine primining injection, tail-pinch and CNO infusion into the LH in hM4Di-expressed rats (n=6).

**Figure 6.**
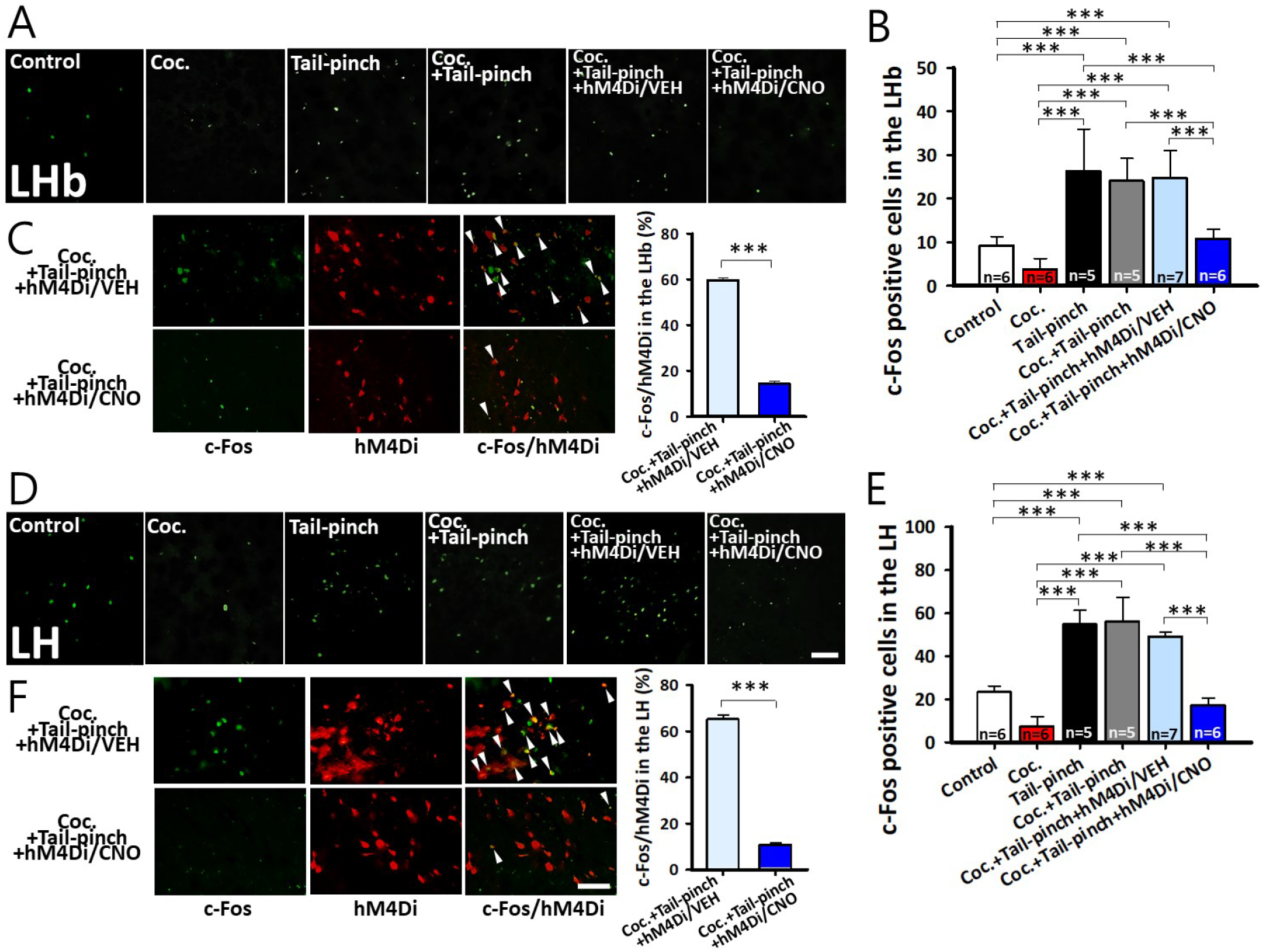
Effects of tail-pinch and chemogenetic silencer on c-Fos expression in the LH and the LHb neurons. **(A)** Representative images of c-Fos expression in the LHb in Control (n=6), Coc. (n=6), Tail-pinch (n=5), Coc.+Tail-pinch (n=5), Coc.+Tail-pinch+hM4Di/VEH (n=7), and Coc.+Tail-pinch+hM4Di/CNO (n=6) groups. **(B)** The number of the c-Fos positive neurons in the LHb in each group. ***p<0.001. **(C)** Representative images of c-Fos immunoreactivity, hM4Di viral expression, and c-Fos expression in the hM4Di-infected neurons (indicated by arrowheads) in the LHb of Coc.+Tail-pinch+hM4Di/VEH and Coc.+Tail-pinch+hM4Di/CNO groups (left) and ratios of the c-Fos positive hM4Di-infected neurons to hM4Di-infected neurons in the LHb (right; ***p<0.001, Coc.+Tail-pinch+hM4Di/VEH vs. Coc.+Tail-pinch+hM4Di/CNO). **(D)** Representative images of c-Fos expression in the LH in Control (n=6), Coc. (n=6), Tail-pinch (n=5), Coc.+Tail-pinch (n=5), Coc.+Tail-pinch+hM4Di/VEH (n=7), and Coc.+Tail-pinch+hM4Di/CNO (n=6) groups. Bar=4O μm. **(E)** The number of c-Fos positive neurons in the LH in each group. ***p<0.001. **(F)** Representative images of c-Fos immunoreactivity, hM4Di viral expression, and c-Fos expression in the hM4Di-infected neurons (indicated by arrowheads) in the LH of the Coc.+Tail-pinch+hM4Di/VEH and Coc.+Tail-pinch+hM4Di/CNO groups (left) and ratios of the c-Fos positive hM4Di-infected neurons to all the hM4Di-infected neurons in the LH (right; ***p<0.001, Coc.+Tail-pinch+hM4Di/VEH vs. Coc.+Tail-pinch+hM4Di/CNO). Bar=4O μm.

### Elevation of c-Fos Expression by Tail-Pinch and its Reversal by Chemogenetic Silencing of the LH-LHb Pathway

Finally, neuronal activities of the LH and the LHb following cocaine, tail-pinch, and/or either CNO or aCSF were evaluated by immunohistochemistry for c-Fos. Tail-pinch increased c-Fos expression in the LH and the LHb in cocaine naïve rats (Tail-pinch group) and the cocaine-injected rats (Coc.+Tail-pinch group), compared to the Control group and the Coc. group (Fig. 5A, 5B, 5D, and 5E). In the rats with LH-LHb:hM4Di, the increased c-Fos expression by tail-pinch was significantly inhibited by CNO administration (Coc.+Tail-pinch+hM4Di/CNO group), but not by aCSF administration (Coc.+Tail-pinch+hM4Di/VEH group; one-way ANOVA for LHb or LH group is F_(5,25)_=19.763 and F_(5,31)_=77.686, respectively). Furthermore, while the ratios of the c-Fos-expressing hM4Di-infected neurons to all the hM4Di-infected neurons were 59.44 ± 1.33% and 65.04±2.10% in the LHb and the LH in the Coc.+Tail-pinch+hM4Di/VEH group, the ratios of that were 14.16 ± 1.25% and 10.66 ± 0.97% in the LHb and the LH in the Coc.+Tail-pinch+hM4Di/CNO group (Fig. 5C and 5F), indicating that, compared to aCSF, CNO infusion significantly decreases c-Fos expression induced by tail-pinch in the hM4Di-infected LH and LHb neurons (unpaired *t*-test).

## Discussion

In the present study, tail-pinch excited LH and LHb neurons, which was blocked by chemical lesion of the LH. Tail-pinch decreased DA levels in the NAc ventrolateral shell, which was disrupted by chemical lesion of the LH. In addition, tail-pinch attenuated cocaine-induced locomotor activity, emission of 50-kHz USVs, development of cocaine-taking behavior, and reinstatement of cocaine-seeking behavior, which was reversed by chemogenetic silencing of the LH-LHb pathway. Tail-pinch increased c-Fos expression of the LH and the LHb neurons, which was inhibited by chemogenetic silencing of the LH-LHb pathway with CNO. These results suggest that nociceptive stimulation suppresses mesolimbic DA system and cocaine-reinforcing effects through the LH-LHb pathway.

The LH projects glutamatergic inputs to the LHb area, which afferents from the LH directly connect to the VTA-projecting LHb neurons (LHb→VTA neurons) [10]. Previous reports have revealed that an anterograde tracer injected into the LH was expressed in the LHb and strongly overlaid with immunocytochemical localization of vesicular-glutamate transporter 2 (VGluT2) in the LHb [10]. The anterogradely traced LH axons overlapped with the LHb neurons which were retrogradely traced by the VTA [10]. Furthermore, whole-brain mapping of neurons projecting to the LHb revealed that the LH is the most prominent input region to the LHb [18]. In the present study, excitation of the LHb by tail-pinch was blocked by the LH lesion and a retrograde viral vector encoding hM4Di which was injected into the LHb was found in the LH area. It suggests that the LHb is directly innervated by the LH and the LH-LHb circuit conveys nociception.

Previous studies revealed that the LH is involved in drug-taking behaviors, reinstatement of drug-seeking behavior, and drug-induced synaptic plasticity [19, 20]. Blacktop and Sorg (2019) reported that degradation of LH structures inhibits cocaine cue-induced reinstatement of drug-seeking behavior in rats [20]. The LHb also plays a critical role in drug-induced craving and aversion [12]. The LH-LHb pathway encodes negative valence and controls motivational behaviors [16, 21]. For example, Lecca et al. reported that chemogenetic silencing of the LH-LHb pathway disrupts escape behaviors against a compartment paired with electric foot shocks and against the abrupt presentation of shadow mimicking an attack of predators [21]. In our findings, both LH and LHb neurons were excited by tail-pinch and chemogenetic silencing of the LH-LHb pathway alleviated inhibitory effects of tail-pinch on cocaine-enhanced locomotor activity and reinstatement of cocaine-seeking behavior, suggesting that nociceptive stimulation inhibits cocaine addictive behaviors through activation of the LH-LHb pathway.

Previous studies have suggested that the LHb is involved in nociceptive processing [9, 22]. It was reported that the LHb responds to noxious, but not to non-noxious stimuli [23]. Consistent with previous studies, our *in vivo* electrophysiological data showed that firing rates of the LHb neurons were evoked by noxious tail-pinch, but not by non-noxious stimuli such as brush and light pressure. We further confirmed excitation of the LHb neurons in response to tail-pinch by using *in vivo* calcium imaging. The LH has also been reported to be critically involved in nociceptive processing [14, 15]. In our study, *in vivo* electrophysiological and calcium imaging data supported that the LH and the LHb neurons are activated by nociception. Furthermore, the tail-pinch-induced activation of the LHb neurons was completely blocked by chemical lesion of the LH. It suggests that nociceptive signals of tail-pinch are conveyed to the LHb via the LH.

Cumulative evidence has suggested that the reward system links to external somatosensory system [24, 25]. Somatosensory stimuli such as noxious stimulation influence the activity of DAergic neurons in the reward system [24]. Activation of cervical spine mechanoreceptors modulates firing of VTA GABA neurons and dopamine release [25]. Furthermore, we and others have shown that somatosensory stimuli reduce drug-induced craving behaviors through modulating the mesolimbic DA systems in rats [26, 27]. Application of acupuncture, widely accepted as a form of peripheral sensory stimulation [28], decreases DA release in the NAc by activating GABA interneurons in the VTA and thus suppresses addiction-related behaviors caused by drugs such as cocaine, morphine, and ethanol [26, 27]. How the somatosensory stimulation affects mesolimbic DA systems is largely unknown. In the present study, tail-pinch reduced the NAc DA release, which was ablated by the LH lesion. Tail-pinch also attenuated cocaine-enhanced locomotor activity and 50-kHz USVs, known to be associated with an increase of the NAc DA level [29], which was inhibited by chemogenetic silencing of the LH-LHb pathway. These findings suggest that somatosensory stimulation such as tail-pinch suppresses mesolimbic DA system via the LH-LHb pathway.

Nociceptive stimulation affects the mesolimbic DA system, although there is a controversy whether the mesolimbic DA release is reduced or increased by nociceptive stimuli. A large body of studies has reported that nociception decreases mesolimbic DA release [30, 31], while other studies have shown opposite results, reporting increase of the mesolimbic DA release by nociceptive stimulation [32, 33]. Interestingly, new insights and attempts were emerged for subtyping the NAc shell according to neuroanatomical or functional features, reporting that the subtypes of the NAc would explain the opposing functions of nociceptive stimuli on the mesolimbic DA release [34, 35]. According to a previous publication, tail-pinch decreased DA release and the activity of DAergic fibers in the ventrolateral part of the NAc shell [34]. On the contrary, tail-pinch increases DA release and the activity of DAergic fibers in the ventromedial part of the NAc shell [34]. de Jong et al. reported that an electric foot shock suppresses the activity of DAergic fibers in the lateral part of the NAc shell whereas it enhances the activity in the ventromedial part of the NAc shell [35]. The present study recorded DA efflux in the ventrolateral part of the NAc shell using the FSCV and found that tail-pinch significantly decreases the mesolimbic DA release. Thus, we assumed that discrepancy in the NAc DA release by nociceptive stimulation in previous studies might be due to different DA recording sites in the NAc.

In conclusion, our findings suggest that the LH-LHb pathway plays an important role in transmitting nociceptive inputs to the mesolimbic DA system and thus inhibiting the cocaine-taking/seeking behaviors.

## Materials and methods

### Animals

All experiments were performed with male Sprague-Dawley rats weighing 250-320 g (Hyochang, Seoul, Korea). Rats had free access to food and water and were kept under the room conditions of a 12-hour light-dark cycle, constant temperature (24 ± 1 °C), and 50% humidity. All procedures were approved by the Institutional Animal Care and Use Committee at Daegu Haany University (DHU2018-824) and conducted according to National Institutes of Health Guide for the Care and Use of Laboratory Animals.

### Chemicals, Reagents, and Antibodies

Cocaine hydrochloride (Macfarlan Smith Ltd, Edinburgh, UK), ibotenic acid (5 μg/μL in saline; Sigma-Aldrich, St. Louis, MO, USA), AAV-hSyn1-GCaMP6s-P2A-nls-dTomato (serotype AAV1, viral titer ≥ 5×10^12^ vg/mL; Addgene, Watertown, MA, USA), pAAV-hSyn-hM4D(Gi)-mCherry (serotype AAV retrograde, viral titer ≥ 7×10^12^ vg/mL; Addgene), and Clozapine N-oxide (CNO) dihydrochloride (Tocris Bioscience, Bristol, UK) were used. Anti-c-Fos antibody (ab190289; Abcam Biotechnology, Cambridge, UK), anti-rabbit IgG antibody (AlexaFluor488, A21206; Thermo Fisher Scientific, Waltham, MA, USA), and Vectashield antifade mounting medium with DAPI (H-1200; Vector Laboratories, Burlingame, CA, USA) were used for the immunohistochemistry.

### Tail-Pinch Stimulation

Tail-pinch stimulation was conducted as described previously with some modifications [36]. Binder clips (19 mm; WHASHIN, Paju, Korea) were used for pinching the tails with a pressing force of 1.0~1.2 kg. Prior to experiments, the pressing force of the binder clips was further ensured by using a force sensor (SW-02; CAS, Beijing, China). The stimulation was applied about 10-20 mm apart from the tips of the tails.

### Chemical Lesion of the LH

As performed in our previous study [37], ibotenic acid (5 μg/μL) was injected 7 days prior to experiments for *in vivo* extracellular single-unit recordings, calcium imaging, and FSCV In brief, rats were anesthetized by intraperitoneal injection (i.p.) with pentobarbital sodium (50 mg/kg) and placed in a stereotaxic frame, and two holes were drilled in the skull to access to the LH (anterior, −2.6 mm; lateral, ±1.7 mm; deep, −8.3 mm). Ibotenic acid or saline was injected into the LH at a rate of 0.25 μL/min for 2 min by using a 26-gauge Hamilton syringe (Reno, NV, USA) and a microinjection pump (Pump 22; Harvard Apparatus, Holliston, MA, USA). The syringe was left in place for at least 5 min to prevent reflux after injection.

### In vivo Extracellular Single-Unit Recording

Rats were anesthetized by urethane (1.5 g/kg, i.p.) and a carbon-filament glass micro-electrode (0.4-1.2 MΩ, Carbostar-1; Kation Scientific, Minneapolis, MN, USA) was stereotaxically inserted into the LH (anterior, −2.6 mm; lateral, ±1.7 mm; deep, −8.3 mm) or the LHb (anterior, −3.5 mm; lateral, ±0.7 mm; deep, −4.9 mm). Single-unit activity was amplified and filtered at 0.1-10 kHz (ISO-80; World Precision Instruments, Sarasota, FL, USA) and then noise was binned from the valid single-unit activity at intervals of 1 sec. The single-unit activities were recorded and analyzed using a CED 1401 Micro3 device and Spike2 software (Cambridge Electronic Design, Cambridge, UK). After recording a stable baseline for at least 10 min, the rats were given brush, light pressure, or tail-pinch for each 10 sec and recorded for a further 10 min.

### In vivo Microendoscopic Calcium Imaging

A calcium sensor GCaMP6s (AAV-hSyn1-GCaMP6s-P2A-nls-dTomato) was injected at a rate of 0.25 μL/min for 2 min (0.5 μL per side) into the left or the right side of the LH (anterior, −2.6 mm; lateral, ±1.7 mm; deep, −8.3 mm) or the LHb (anterior, −3.5 mm; lateral, ±0.7 mm; deep, −4.9 mm) in the rats anesthetized with pentobarbital sodium. An imaging cannula with a 500 μm diameter gradient index (GRIN) lens (Doric Lenses, Quebec, Canada) was then placed into the LH or the LHb and anchored to the skull using dental cement and stainless steel screws. Four to 6 weeks after the surgery, the rats were connected to the microscope body via the imaging cannulas implanted on the heads and then calcium imaging process was conducted. The fluorescent calcium transients were recorded and processed via Doric Neuroscience Studio software (10 frames per second, and 20 ± 5% of illumination power, v.5.2.2.3; Doric lenses). After recording basal calcium activity for 10 sec, the rats were given tail-pinch for 10 sec and recordings were continued for a further 10 sec. The background fluorescence was eliminated from the motion-corrected images. Regions of interests (ROIs) were determined using the algorithm, principal component analysis (PCA)/independent component analysis (ICA) that extracted the distinctive cellular signals from imaging data sets. Finally, the relative fluorescence change (ΔF/F_0_) of each ROIs was calculated with F_0_ (baseline fluorescence) corresponding to the temporal average of fluorescence intensity (ΔF).

### In vivo Fast-Scan Cyclic Voltammetry (FSCV) for Monitoring DA Release

Electrically evoked-DA release in the NAc ventrolateral shell was measured by *in vivo* FSCV, as performed in our previous study [38]. A custom-made carbon fiber electrode (CFE; 7 μm diameter, 200 μm length of exposed tip) was used. The electrode potential was scanned with a triangular waveform from –0.4 to +1.3 V and back to –0.4 V versus Ag/AgCl at a scan rate of 400 V per sec. Cyclic voltammograms were recorded every 100 msec by ChemClamp voltage clamp amplifier (Dagan Corporation, Minneapolis, MN, USA). Recording and analyzing were performed using a LabVIEW-based (National Instruments, Austin, TX, USA) customized Demon voltammetry software. Under urethane anesthesia (1.5 g/kg, i.p.), bipolar stainless steel electrode and CFE were stereotaxically placed into the medial forebrain bundle (MFB; anterior, −2.5 mm; lateral, ±1.9 mm; deep, −8.0~-8.5 mm) and the NAc ventrolateral shell (anterior, +1.6 mm; lateral, ±1.9 mm; deep, −8.0~-8.5 mm). The MFB was stimulated with 60 monophasic pulses at 60 Hz (4 msec pulse width) every 2 min. After a stable baseline was established (less than 10% variability in peak heights of 5 consecutive collections), the changes of DA release in the NAc ventrolateral shell following tail-pinch were monitored for a further 30 min. To identify the recording sites, the rats were sacrificed at the end of the experiments and perfused with 4% formaldehyde. The brains were taken out, stored in 30% sucrose solution and cryo-sectioned coronally at a thickness of 30 μm. The slices were stained with toluidine blue and observed under a light microscope (Microscopesmall, Guangxi, China).

### Chemogenetics and Cannula Implantation

Under pentobarbital anesthesia (50 mg/kg, i.p.), a retrograde viral vector encoding an inhibitory designer receptors exclusively activated by designer drugs (DREADD), hM4Di, was bilaterally injected at a rate of 0.25 μL/min for 2 min (0.5 μL per side) into the LHb by using a 26-gauge Hamilton syringe mounted on a microinjection pump. Four weeks after the viral injection, guide cannulas (26-gauge; Plastics One, Roanoke, VA, USA) were implanted bilaterally into the LH (anterior, −2.6 mm; lateral, ±1.7 mm; deep, −7.3 mm) to locally infuse the DREADD agonist, CNO or artificial cerebrospinal fluid (aCSF). A week after the surgery, experiments for locomotor activity and 50-kHz ultrasonic vocalizations (USVs) were conducted. Internal cannulas (33-gauge; Plastics One) which were protruded beyond the length of guide cannulas by 1 mm were inserted into the guide cannulas implanted on the heads. CNO was dissolved in aCSF to 1 mM, as previously described [39]. CNO or vehicle (aCSF) was intracranially infused at a rate of 0.15 μL/min for 2 min through the internal cannulas connected to a 26-gauge Hamilton syringe and a microinjection pump by polyethylene tubing and an additional 1 min was allowed for further diffusion.

### Measurement of 50-kHz USVs and Locomotor Activity

50-kHz USVs and locomotor activity were recorded simultaneously in customized sound-attenuating chambers. The chamber consisted of two boxes to minimize exterior noise (inside box: 60×42×42 cm, outside box: 68×50×51 cm). A condenser ultrasonic microphone (Ultramic 250K; Dodotronic, Castel Gandolfo, Italy) and a digital camera were positioned at the center of the ceiling of the chamber. As performed in our laboratory [29], 50-kHz USVs were recorded using the ultrasonic microphone with UltraSoundGate 416H data acquisition device (Avisoft Bioacoustics, Glienicke, Germany). Ultrasonic vocal signals were band-filtered between 30- and 80-kHz for the 50-kHz USVs and analyzed using Avisoft-SASLab Pro (version 4.2; Avisoft Bioacoustics). Locomotor activity was measured with a video-tracking system (Ethovision XT; Noldus Information Technology BV, Wageningen, Netherlands). Rats were habituated for 30 min in the chambers. After recording basal USVs and basal activity for 30 min, the rats received cocaine (15 mg/kg in saline, i.p.), tail-pinch, and/or either CNO (1 mM) or aCSF through the implanted guide cannulas. The recordings were continued for a further 60 min.

### Cocaine Self-Administration, Extinction, and Reinstatement Procedures

Food training and cocaine self-administration were performed in operant chambers equipped with the active and inactive levers (Med Associates, St. Albans, VT, USA) as described previously with slight modifications [40]. Initially, rats were food-restricted with 16 g of lab chow per day and trained to press the active lever to gain 45 mg food pellets (Bio-Serve, Frenchtown, NJ, USA). Rats that achieved criterion for food responding (100 food pellets for three consecutive days) were chosen for the cocaine self-administration procedure and surgically implanted with chronically indwelling intravenous catheters under pentobarbital anesthesia. After recovery of at least 7 days, the rats were trained to self-administer cocaine intravenously by pressing the active lever under a fixed-ratio schedule (FR 1) in a daily 2-h session. Once the active lever was pressed, a 0.5 mg/kg/0.1 mL cocaine was infused over 5 sec. When the intravenous cocaine infusion was initiated, the house light was extinguished for 20 sec and a cue light located above the active lever was illuminated for 5 sec, concomitant with a 15-sec time-out period. During the time-out period, the active lever responses were recorded in an automated counting program (Schedule Manager; Med Associates), but no cocaine infusion was made. Pressing the inactive lever produced no scheduled responses but signals were recorded in the program (Schedule Manager; Med Associates). After 10 sessions of the cocaine self-administration training, the rats underwent 7 sessions of the extinction task during which saline was delivered to the rats without an illumination of the cue light when they had pressed the active lever. After the extinction task, the rats received a single priming intravenous injection of 0.5 mg/kg cocaine and the experiment was performed with the same experimental conditions of the extinction task.

### Immunohistochemistry

The immunohistochemistry for c-Fos was carried out as described previously [41]. Forty min after tail-pinch, cocaine injection, and/or either CNO or aCSF infusion, the rats were perfused with 4% formaldehyde. The brains were taken out, post-fixed, cryo-protected and cryo-sectioned into 30 μm-thick. The brain slices were incubated with anti-c-Fos rabbit polyclonal antibodies (1:500), followed by donkey anti-rabbit IgG antibodies (Alexa Fluor 488, 1:500). The slices were then mounted on gelatin-coated slides, photographed and examined under a confocal laser scanning microscope (LSM700; Carl Zeiss, Oberkochen, Germany). The number of c-Fos positive cells in the LHb or the LH was blindly counted and 5-7 slices per animal were analyzed.

### Statistical Analysis

Data were presented as the mean ± standard error of the mean (SEM) and analyzed by one- or two-way measurement analysis of variance (ANOVA), one- or two-way repeated ANOVA, followed by post hoc testing using the Tukey method, unpaired *t*-test or paired *t*-test, where appropriate. Values of p<0.05 were regarded as statistically significant.

## Acknowledgements

HYK conceived and designed research. SML and YF performed the research. SML, YF and HYK analyzed the data and drafted the manuscript. HYK was responsible for the overall direction of the project and for the editing of the manuscript. SCS and KBB provided feedback on the project. All authors read and approved the final manuscript. This research was supported by the National Research Foundation of Korea (NRF) grant funded by the Korea government (MSIT) (No. 2018R1A5A2025272, 2019R1A2C1002555, and 2019R1A6A3A13090969) and the Korea Institute of Oriental Medicine (KIOM) (KSN1812181, KSN2013210).

